# The parathyroid hormone-dependent activation of osteoblasts enhances hematopoietic stem cell migration and reduces their engraftment abilities

**DOI:** 10.1101/2021.03.04.433901

**Authors:** Yasmine Even, Lin Yi, Chih-Kai Chang, Fabio M.V. Rossi

## Abstract

Hematopoietic stem cells (HSCs) in the bone marrow (BM) reside in HSC niches ensuring their maintenance. The HSC niche is made up of perivascular and trabecular cells including osteoblasts whose role on HSCs remains to be clearly defined. Increased numbers of osteoblasts have been observed in the CL2 transgenic mouse expressing a constitutively activated form of the parathyroid hormone (PTH)/PTH-related peptide receptor. This mouse model mimicking PTH anabolic effect has also been described to exhibit increased numbers of the BM stem/progenitor population. Furthermore, PTH is known to induce BM stem/progenitor cell migration into blood circulation. However PTH role on long-term repopulating HSCs (LT-HSCs) is incompletely known. Here we show that CL2 BM contains a regular proportion of LT-HSCs, suggesting that osteoblasts may not be a determinant of LT-HSC numbers but act mainly on more mature progenitors. We further show increased LT-HSC migration in CL2 mice correlated with higher granulocyte colony-stimulating factor (G-CSF) serum levels, supporting the idea that PTH can enhance the migration of LT-HSCs. Finally, we found a defect in the ability of CL2 BM HSCs to reconstitute irradiated BM suggesting that PTH activation of osteoblasts negatively influences abilities of HSC population to engraft and reconstitute irradiated BM. In summary, our study highlights new insights into the role of the PTH-dependent activation of osteoblasts on LT-HSC migration and their BM repopulation abilities. Our findings will be useful to improve treatments on hematological disorders, especially therapies involving HSC harvest and transplantation.

## Introduction

In the bone marrow (BM), hematopoietic stem cells (HSCs) require a specific microenvironment regulating their maintenance and proliferation and called the HSC niche. The role of niche associated cells and their interactions with HSCs are complex and remain incompletely known [1]. The first BM HSC niche described, located next to the osteoblast composed endosteum was called the endosteal niche [2–3]. The second HSC niche defined, the perivascular niche, was located next to vessels in BM and was associated with mesenchymal stromal and endothelial cells [4–5]. More recent studies have specified the location of BM HSC niches by showing that HSCs are found near sinusoids as well as in the endosteal area next to arterioles [6–10]. However while it is now well established that BM HSC niches are located in perivascular areas, the endosteal-vascular niche is more controversial [11]. Indeed, deep confocal imaging studies have shown that BM HSCs are mainly located next to sinusoidal blood vessels [8]. Recent reviews highlight the remodeling BM throughout lifespan especially BM stroma and vascular layout, which would influence HSC niche cell composition and localization [12–13]. The BM HSC niche was suggested to be close to the endosteum during perinatal period to gradually evolve and become a central sinusoid HSC niche in adulthood [13].

The role of osteoblastic cells on HSCs and whether they play a role as part of the niche remains unclear. *In vitro* studies have proposed osteoblasts as being an important part of the endosteal niche by secreting factors essential for stem cell maintenance [14]. Two *in vivo* studies showed that transgenic animals with increased osteoblast activity through stimulation of the parathyroid hormone/parathyroid hormone-related protein receptor (PPR) or inactivation of the bone morphogenic protein receptor-1a, exhibited a correlated increase in the number of BM stem/progenitor cells [2–3]. Moreover, a recent study demonstrated that the loss of endosteal BM occurring in acute myeloid leukemias was correlated with HSC and osteoblastic cell depletion [15]. However, Visnjic et al. reported that conditional ablation of osteoblasts did not induce any severe loss in the BM HSC pool size, but rather promotes a high loss of more differentiated cells, especially B cell progenitors [16]. Furthermore, a study using strontium as a treatment to expand osteoblasts, showed that it was not correlated with HSC increase [17]. And more recently, Calvi et al. showed that PPR dependent expansion of osteocytes and mature osteoblasts did not increase BM HSCs [18]. Therefore, the role of osteoblasts on the BM HSC population remains to be clarified.

It is know well established that osteoblasts can be influenced by the parathyroid hormone (PTH). More precisely, PTH can exert a catabolic or anabolic effect on bone. While a continuous delivery of PTH induces bone loss, intermittent PTH distribution induces an opposite effect. Calvi and collaborators have described the action of PTH on bones mediated by the PPR. In fact, the transgenic mouse (CL2 mouse) expressing the constitutively activated form of PPR in its osteoblasts (Col2.3 promoter) exhibits higher numbers of osteoblasts and increased trabecular bone [19]. This study describes that constitutive PPR activation in osteoblastic cells mimics the anabolic effect of PTH, inducing bone increase by enhancing osteoblast function. In CL2 mice, the higher numbers of osteoblasts in the trabecular compartment and at the endosteal surface of cortical bone is due to a combined increased proliferation and decreased apoptosis. Very few studies have been performed on the role of PTH on HSCs. However, it is known that PTH induces the mobilization of hematopoietic progenitor cells without depleting the BM [20]. Furthermore, Adams et al. have shown that PTH treatment increases HSC mobilization induced by granulocyte colony-stimulating factor (G-CSF) and also protects HSCs from multiple exposure to cytotoxic chemotherapy [21]. However, the role of PTH on the migration and repopulation properties of long-term repopulating HSCs (LT-HSCs) remains poorly known. The PTH molecule has been proposed as new potential mobilizating agent for transplantation therapies. The success of transplantation in treatments for hematological disorders partly depends on the capacity of HSCs to reconstitute the BM. Therefore, a better understanding of PTH action on HSC properties will be useful to improve hematological disorder treatments involving HSC mobilization and BM repopulation and to develop new therapeutic strategies.

In the present study, we show that the osteoblast increase induced by PPR activation in CL2 mice correlates with an increase of BM progenitor cells whereas the proportion of LT-HSCs remains unchanged. We also found higher LT-HSC migration in CL2 mice. Finally, we show that CL2 BM HSCs have impaired ability to reconstitute irradiated BM.

## Materials and methods

### Animals

All mice were bred in-house and maintained in a pathogen-free environment. CL2 mice (also called Col1-caPPR mice) have been described in [19]. Briefly, the constitutively active PPR-H223R is under the control of the mouse 2.3 Col1A1 promoter (bone specific). F1 from CL2-FVB/N male (kind gift from E. Schipani) and heterozygous-GFP^+^ Ly5.2 C57BL/6 female, expressing GFP ubiquitously from the cytomegalovirus enhancer-chicken β-actin hybrid promoter [22], were used for all the experiments (WT and CL2 littermates). F1 from FVB/N male and C57BL/6 Ly5.1 female mice, 8-12 weeks of age, were used as recipients. All experiments were approved by the University of British Columbia Animal care committee (A05-0351, A06-0185).

### Antibodies and staining reagent for flow cytometry analyses

Phycoerythrin (PE) conjugated anti-c-kit and Peridinin Chlorophyll Protein Complex (PerCP) or allophycocyanin (APC) conjugated anti-CD45 antibodies were purchased from BD Biosciences Pharmingen; anti-CD3, anti-B220, anti-Mac-1, anti-Gr-1 & anti-TER119 antibodies conjugated to PE-Cyanin7 (PE-Cy7) or to biotin, anti-CD45.2 and anti-Sca-1 antibodies conjugated to APC and streptavidin conjugated to eFluor450 or conjugated to PE-Cy7 were purchased from eBioscience. Anti-CD150 antibodies conjugated to fluorescein isothiocyanate (FITC), to biotin or to PE-Cy7, anti-CD201 antibodies conjugated to PE and anti-CD48 antibodies conjugated to APC or PE-Cy5 were purchased from Biolegend (San Diego, CA). To stain dead cells, 4′,6-diamidino-2-phenylindole (DAPI) was purchased from Sigma-Aldrich. For blood recipient analyses: PE conjugated anti-CD3 was purchased from BD Biosciences Pharmingen; PE conjugated anti-B220, PE-Cy7 conjugated anti-Gr-1 and anti-Mac1 and APC conjugated anti-CD45.2 were purchased from eBioscience.

### Population analysis and quantification by flow cytometry

We used flow cytometry to quantify the proportion and total numbers of KLS population and HSCs in CL2 and WT BM and spleen. BM was flushed off the femurs and tibiae. After red blood cell lyses, nucleated BM cells were stained for KLS and KLS-SLAM population analysis with the following antibodies: anti-CD3, anti-B220, anti-Ter119, anti-Gr1 and anti-Mac1 conjugated to biotin or conjugated to PE-Cy7 (Lin labelling), anti-C-Kit conjugated to PE, anti-Sca1conjugated to APC, anti-CD150 conjugated to PE-Cy7 or conjugated to FITC and anti-CD48 conjugated to PECy5, followed by a streptavidin-eFluor^®^ 450 staining when biotin-conjugated antibodies were used. For E-SLAM population analysis, staining was realized with the following antibodies: APC-conjugated or PE-Cy5-conjugated anti-CD48, biotin or FITC-conjugated anti-CD150, PE-conjugated anti-CD201, PerCP or APC-conjugated anti-CD45 antibodies and streptavidin-PE-Cy7 when biotin-conjugated anti-CD150 antibody was used. To quantify the total numbers of cell populations, cell counting beads were added to each sample as recommended by the manufacturer (Life Technologies). Data were collected with a FACSCalibur or a LSR II flow cytometer (Becton Dickinson) and analyzed with FlowJo software (TreeStar).

### Parabiosis and separation surgery

We used F1 from CL2-FVB male and heterozygous-GFP^+^ Ly5.2 C57BL/6 female in order to perform parabiosis in syngenic conditions to avoid immune reaction from partners. Pairs of 8- to 12-week-old female littermates were housed together in a single cage for 2 weeks and thereafter were joined in parabiosis. Mice were anesthetized with isoflurane (1.5-2%, inhaled) and joined as described in [23–24]. Briefly, on one side of each mouse, skin was incised, olecranon and knee joints were attached by suture and tie and mouse skins were sutured by staples. Parabionts were surgically separated 6 weeks after parabiosis surgery by a reversal of the above procedure.

### BM and spleen competitive repopulation assays

Lethally irradiated (9.5 Gy) 8- to 12-week-old mice (CD45.1) received intravenously 1×10^7^ total BM cells or 5×10^7^ splenocytes from 8-week-old F1 FVB-C57BL/6 donors at a 1:1 ratio of the following donor strains: WT (CD45.1/CD45.2) versus WT or CL2 (CD45.1/CD45.2/GFP^+^) sex-matched littermates. Sixteen weeks post-transplantation, recipients’ blood was collected and single-cell suspensions were stained for myeloid, lymphoid and CD45.2 markers. The relative contribution of BM reconstitution from donors was assessed by quantifying the proportion of GFP^+^ cells among the CD45.2^+^ population by flow cytometry on a FACSCalibur. For each group, donor cells were injected into at least 3 recipient mice.

### BM transplantation of separated parabiont

BM transplantation was performed to evaluate the ability of HSCs to self-renew after migration. The BM donor cells were harvested from parabionts 16 weeks after separation and 5.10^6^ nucleated BM cells were transplanted into lethally irradiated WT congenic recipients (Ly5.1). BM from each donor was transplanted into at least 3 recipients. Sixteen weeks post-transplantation, peripheral blood sample was collected from recipients and erythrocytes were lysed. Nucleated cells were stained using antibodies to CD45.2 and to myeloid (Mac-1, Gr-1) or lymphoid (CD3, B220) lineage markers and analyzed by flow cytometry on a FACSCalibur.

### Competitive repopulating unit (CRU) assay

For *in vivo* quantification of long-term HSCs in WT and CL2 mice, a CRU assay was performed [25]. Briefly, different dilutions (5×10^3^ to 1.25×10^5^) of BM cells from WT or CL2 mice (GFP) were co-injected with 10^5^ competitor WT (Ly5.2) BM cells into lethally irradiated (9.5 Gy) recipients (Ly5.1). More than 16 weeks after transplantation, the level of donor-derived (GFP^+^) myeloid and lymphoid cells in peripheral blood was quantified by flow cytometry. Recipients with more than 2% donor-derived myeloid and lymphoid cells were considered to be repopulated by at least one CRU. CRU frequency was calculated by applying Poisson statistics to the rate of negative recipients at different dilutions using L-Calc software (Stem Cell Technologies).

### Histology and imaging

Spleen from WT and CL2 mice was harvested, fixed, processed and stained using standard procedures (n=4). Haematoxylin & Eosin stained cryostat sections (7μm) of spleen were visualized using a microscope (Zeiss Microimaging) and images were acquired using a charge-coupled device camera (qImaging) and OpenLab4 software (Improvision).

### Measurement of G-CSF concentrations in mouse serum

Blood was harvested from terminally anesthetized 6- to 12-wk-old WT and CL2 mice by cardiac puncture. After clot formation serum was collected. The level of G-SCF in serum samples was measured using an ELISA kit according to the manufacturer’s instructions (RayBiotech).

### Statistical methods

Data are presented as mean ± SE. At least three biological replicates were analyzed, and differences were considered statistically significant by a Student two-tailed t test for a P≤ 0.05. Statistical analyses were performed using Microsoft Excel (Microsoft, Redmond, WA). For CRU assay, Poisson statistics were performed using the L-Calc software (Stem Cell Technologies).

## Results

### CL2 mice exhibit increased KLS population and normal HSC proportion in BM

CL2 mice under the FVB strain have previously been shown to undergo higher proportion of osteoblasts and of hematopoietic stem/progenitor cells [2, 19]. We first measured the proportion of BM stem/progenitor cells present in CL2 and WT mice by performing flow cytometry analyses of the c-Kit^+^Lin^−^Sca1^+^ (KLS) population. In accordance with results in previous studies, a higher proportion of KLS cells was observed in CL2 BM (Fig. 1A upper plots & 1B). We further wanted to assess the proportion of HSCs in CL2 BM. To do so, we first used signaling lymphocyte activation molecule (SLAM) family markers (CD150^+^CD48^−^) within the KLS population. This combination of markers has previously been shown to highly increase the purity of HSC population [4]. To our surprise, we found a similar proportion of BM CD48^−^CD150^+^KLS cells in CL2 and in WT mice (Fig. 1A lower plots & 1C). In order to straighten this result we used the combination of SLAM (CD48^−^ and CD150^+^) and of E-PCR receptor (CD201^+^) markers allowing to gate for a population, called E-SLAM, containing at least ~60% of LT-HSCs [26]. In accordance with our previous results, the proportion of E-SLAM cells was not significantly different in CL2 and WT BM (Fig. 1D & E). In order to more precisely know the frequency of BM HSCs in CL2 mice we realized a competitive repopulating unit (CRU) assay. No significant difference in the proportion of CRUs was observed in CL2 and WT BM (Fig. 1F). All these data show that CL2 mice exhibit higher frequency of the KLS population in BM than WT mice whereas the proportion of CL2 and WT BM HSCs is comparable.

**Figure 1:**
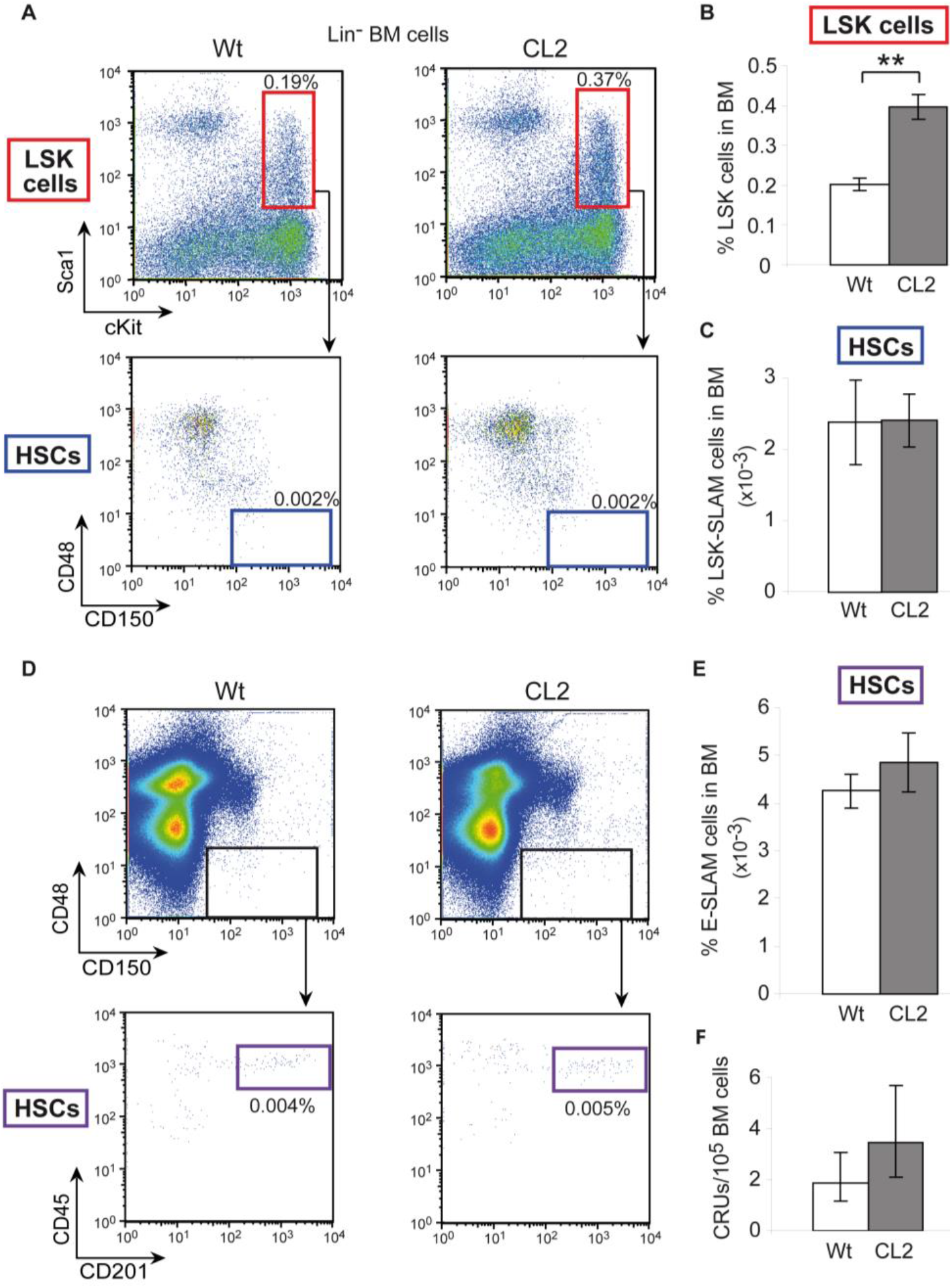
Higher proportion of progenitor cells but same proportion of long-term HSCs in CL2 BM. A. Representative dot plots of Lin^−^ gated BM from WT and CL2 mice. Gated regions designate stem/progenitor cells (KLS cells, Lin^−^ Sca^+^ Kit^+^) on upper plots and HSCs (Lin^−^ Sca^+^ Kit^+^ CD48^−^ CD150^+^) on lower plots. B&C Proportion of stem/progenitor cells (KLS cells, Lin^−^ Sca^+^Kit^+^) (B) and HSCs (Lin^−^Sca^+^Kit^+^CD48^−^CD150^+^) (C) in BM, n=6. D. Representative dot plots of live BM cells (PI^low^) from WT and CL2 mice. The CD48 and CD150 SLAM markers (upper plots) and hematopoietic-(CD45) and CD201 E-PCR-markers (lower plots) were used to isolate HSCs (CD48^−^ CD150^+^ CD45^+^ CD201^+^). E. Proportion of HSCs (CD48^−^ CD150^+^ CD45^+^ CD201^+^) in BM, n=3. F. Proportion of CRUs in WT and CL2 BM. Values indicate the mean percent (± SEM) of at least 3 mice. FlowJo software from TreeStar (Ashland, OR) was used to create plots. **P<0.01.

### CL2 BM contains regular numbers of KLS cells and low numbers of HSCs due to low BM cellularity

As shown by Calvi et al., CL2 mice are smaller and have shorter limbs (Supplementary Fig. S1) however they exhibit increased trabecular bone volume due to higher osteoblast numbers and activity, resulting in remodeling of marrow cavity [19]. In order to know whether these parameters would influence BM-HSC numbers in CL2 mice, we calculated the total numbers of BM cells and of KLS and HSC populations in CL2 bones (femurs and tibias). Our results show 1/3 less BM cells in CL2 than in WT mice (Fig. 2A). Despite the low BM cellularity in CL2 mice, due to higher KLS cell proportion in CL2 BM, the total numbers of KLS cells are similar in CL2 and WT BM (Fig. 2B). In contrast, total numbers of LT-HSCs in BM, are significantly lower in CL2 than in WT bones, when labeled with E-SLAM markers (Fig. 2C, D&E). These data indicate that in comparison to WT, the CL2 mice exhibit lower BM cellularity resulting in lower numbers of BM HSCs.

**Sup. Fig 1:**
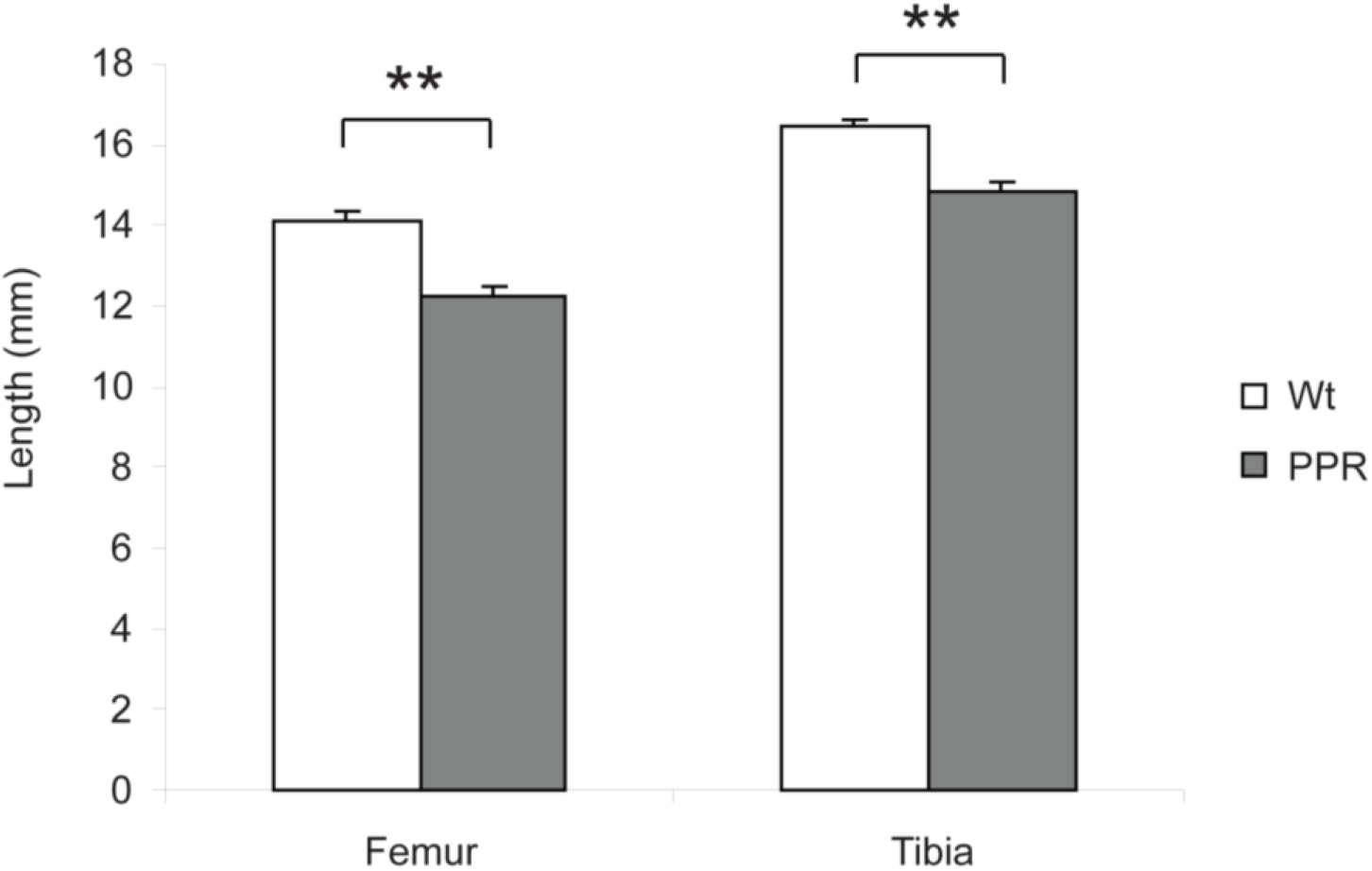
Smaller bone length in CL2 mice. Graph shows the length of femur and tibia in WT and CL2 mice, n=4. **P<0.01.

**Fig 2:**
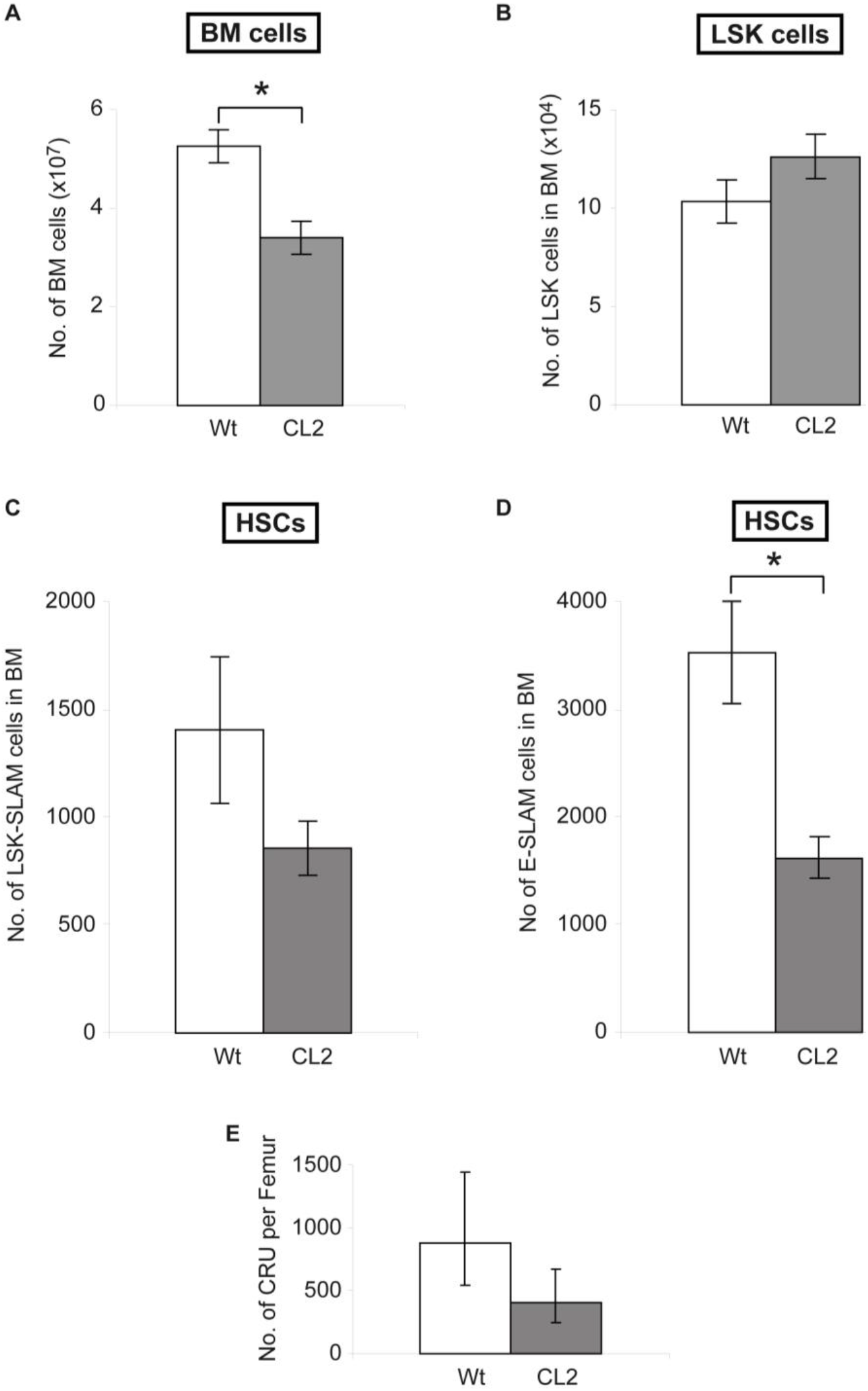
Lower numbers of HSCs in CL2 mice due to lower BM cellularity. Total numbers of BM cells, n=6 (A), KLS cells (stem/progenitor cells), n=6 (B), KLS-SLAM cells (HSCs), n=6 (C) and E-SLAM cells (HSCs), n=3 (D) in WT and CL2 BM (2 femurs + 2 tibias). E. Graph shows numbers of CRUs per femur. Values indicate the mean percent (± SEM) of at least 3 mice. *P<0.05.

### CL2 mice present high numbers of HSCs in spleen without extramedullary hematopoiesis

Because BM cellularity is lower in CL2 mice we investigated whether HSCs would leave BM to seek somewhere else in the body, more likely to the spleen. We first compared the proportion of KLS cells present in CL2 and WT spleen. We found a significantly higher frequency of spleen KLS cells in CL2 than in WT mice (Fig. 3A). We also quantified the proportion of spleen HSCs by using the E-SLAM labeling protocol. We found a proportion of HSCs 4 times higher in CL2 than in WT spleen (Fig. 3B&C). We further quantified the total numbers of spleen HSCs and we found an equivalent HSC number increase in CL2 spleen (Fig. 3D). Moreover, in order to quantify the LT-HSCs, we performed a competitive long-term repopulation assay with CL2 and WT splenocytes. The mutant PPR is not expressed on HSCs due to its collagenI-driven expression [19]. Therefore it should not intrinsically influence the ability of HSCs to reconstitute BM and the participation of HSCs in reconstituting irradiated BM should reflect the proportion of HSCs present within the splenocyte population. At 16 weeks after transplantation, the BM reconstitution ability of HSCs from CL2 and WT spleen was measured by flow cytometry. Consistent with our previous results, the participation of CL2 spleen HSCs in reconstituting recipient BM was much greater than the WT ones (Fig. 3E). This result confirms a very high proportion of LT-HSCs in CL2 spleen. As higher numbers of HSCs in spleen might be due to extramedullary hematopoiesis, we further assessed whether this event was happening in CL2 mice. Extramedullary hematopoiesis is characterized by splenomegaly and the presence of megakaryocytes as well as erythroid and myeloid colonies. None of such colonies were found in CL2 mice neither significant differences in spleen size and cellularity (Fig. 4A, B&C). In absence of extramedullary hematopoiesis, the numbers of HSCs present in spleen often reflect the numbers of HSCs circulating in blood. Therefore, these results suggest high HSC migration into the blood stream of CL2 mice.

**Fig 3:**
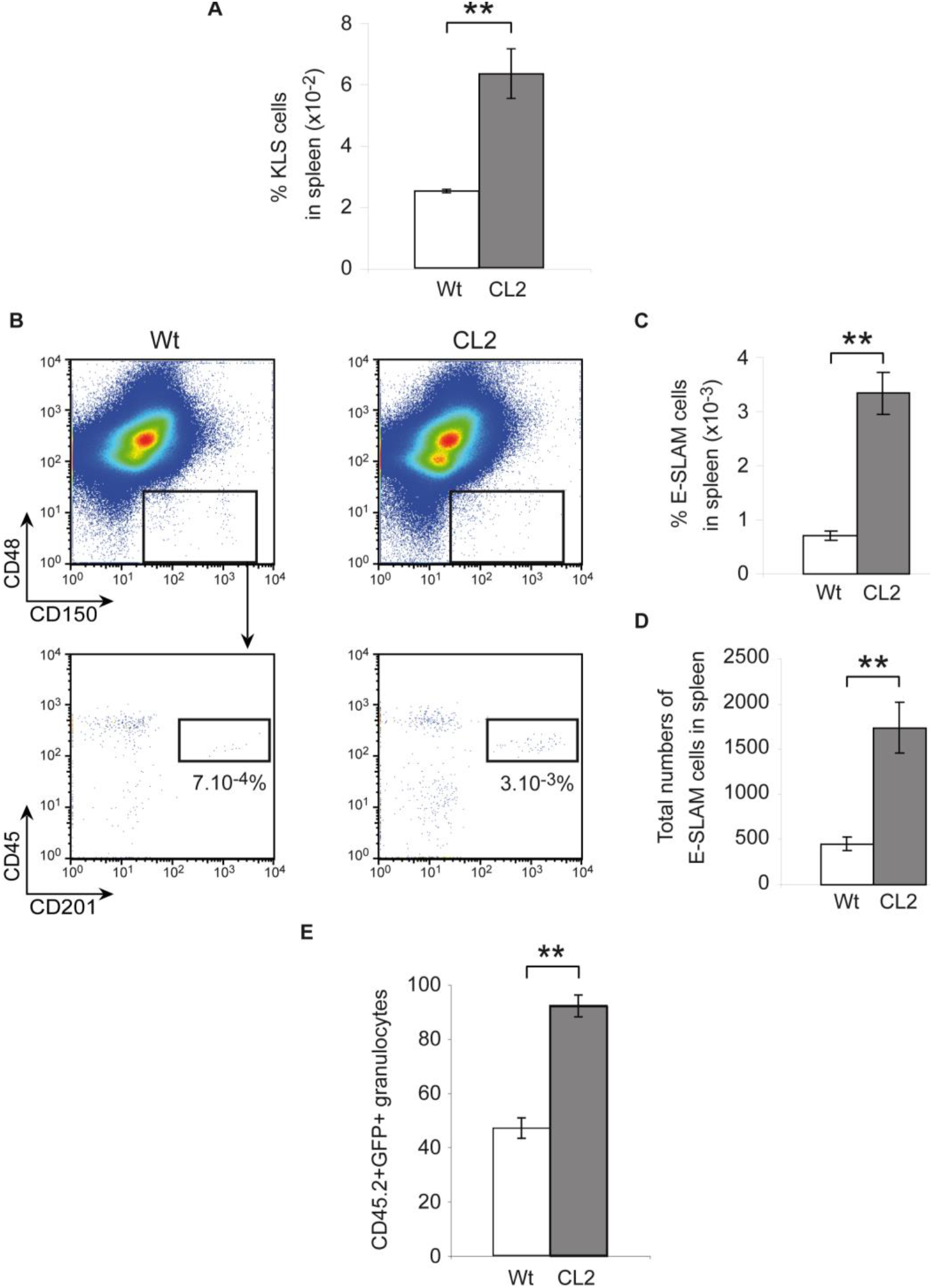
Higher numbers of HSCs in CL2 spleen. A. Graph shows the proportion of KLS cells in WT and CL2 spleen, n=3. B. Representative dot plots of live spenocytes (PI^low^) from WT and CL2 mice. The CD48 and CD150 SLAM markers (upper plots) and hematopoietic-(CD45) and CD201 E-PCR-markers (lower plots) were used to isolate HSCs (CD48^−^CD150^+^CD45^+^CD201^+^). C&D. Proportion (C) and total number (D) of HSCs in WT and CL2 spleen, n=3. E. Graph representative of 3 independent competition assays. **P<0.01.

**Fig 4:**
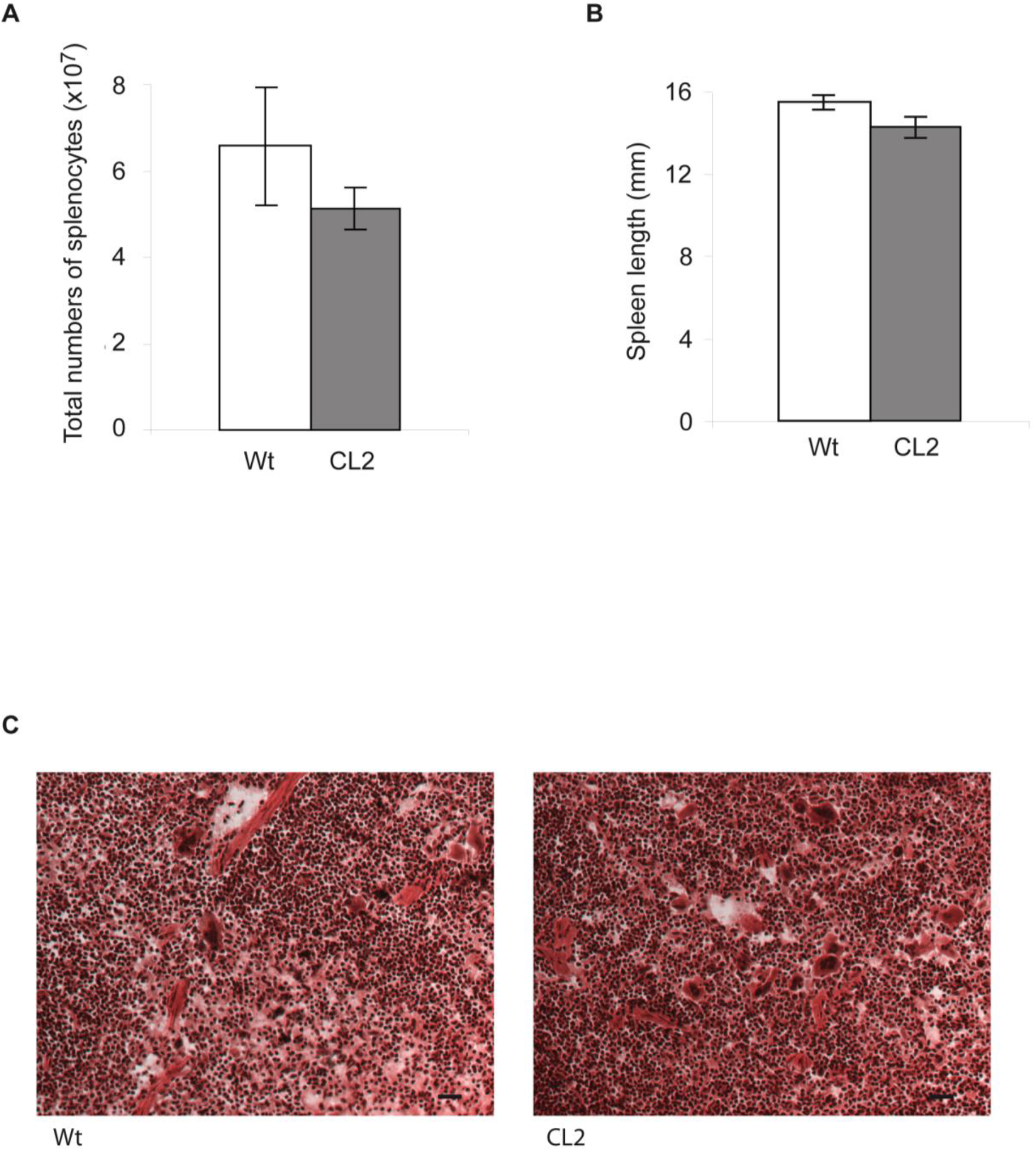
No extramedullary hematopoiesis in CL2 spleen. A. Graph shows total numbers of CL2 and WT splenocytes, n=3. B. Graph shows WT and CL2 spleen length, n=4. C. H&E section of WT and CL2 spleen, magnification x200, scale bar = 50 μm.

### Elevated HSC migration in CL2 mice

In order to measure the frequency of circulating HSCs in CL2 blood stream, we used the parabiosis technique. Two CL2 or 2 WT mice (one mouse GFP and its mouse-partner non-GFP) were joined in parabiosis for 6 weeks. Thereafter parabiosis pairs were separated for 16 weeks before blood and BM harvesting (Fig. 5A). The proportion of HSCs that have migrated into the partner and engrafted its BM was measured by flow cytometry for each separated parabiont. Granulocytes are short term living cells and consequently usually chosen to measure the proportion of LT-HSCs in parabiosis (Fig. 5B). Our analyses show two times more CL2 than WT granulocytes issued from migrated HSCs, indicating that twice as many CL2-HSCs as WT-HSCs have migrated and engrafted the partner’s BM (Fig. 5B&C). This result suggests that in CL2 mice at least two times more HSCs migrate from BM to blood circulation and are thereafter able to re-engraft BM. We further wanted to check the BM repopulation ability of HSCs that have migrated and re-engrafted the partner’s BM. We consequently transplanted the BM of separated parabionts into lethally irradiated Ly 5.1 recipients. Recipient’s blood was analyzed for myeloid and lymphoid populations within donor derived cells. In accordance with our previous result, the proportion of myeloid, B and T cells issued from migrated HSCs were two times higher in recipients transplanted with CL2 parabiont BM than in the ones transplanted with WT parabiont BM (Fig. 5D). These results show that more CL2 LT-HSCs circulate in bloodstream than WT ones. Among the factors known to induce HSC mobilization into blood circulation, G-CSF is one of the most important. Furthermore, PTH injected mice have been shown to exhibit higher G-CSF levels [20]. We therefore measured the level of serum G-CSF in CL2 and WT mice. In agreement with the high frequency of HSC migration in CL2 mice the level of serum G-CSF is also higher in CL2 than in WT mice (Fig. 5E). Overall, these results suggest that the PTH-dependent PPR activation in osteoblasts of CL2 mice induces LT-HSC migration into blood circulation.

**Fig 5:**
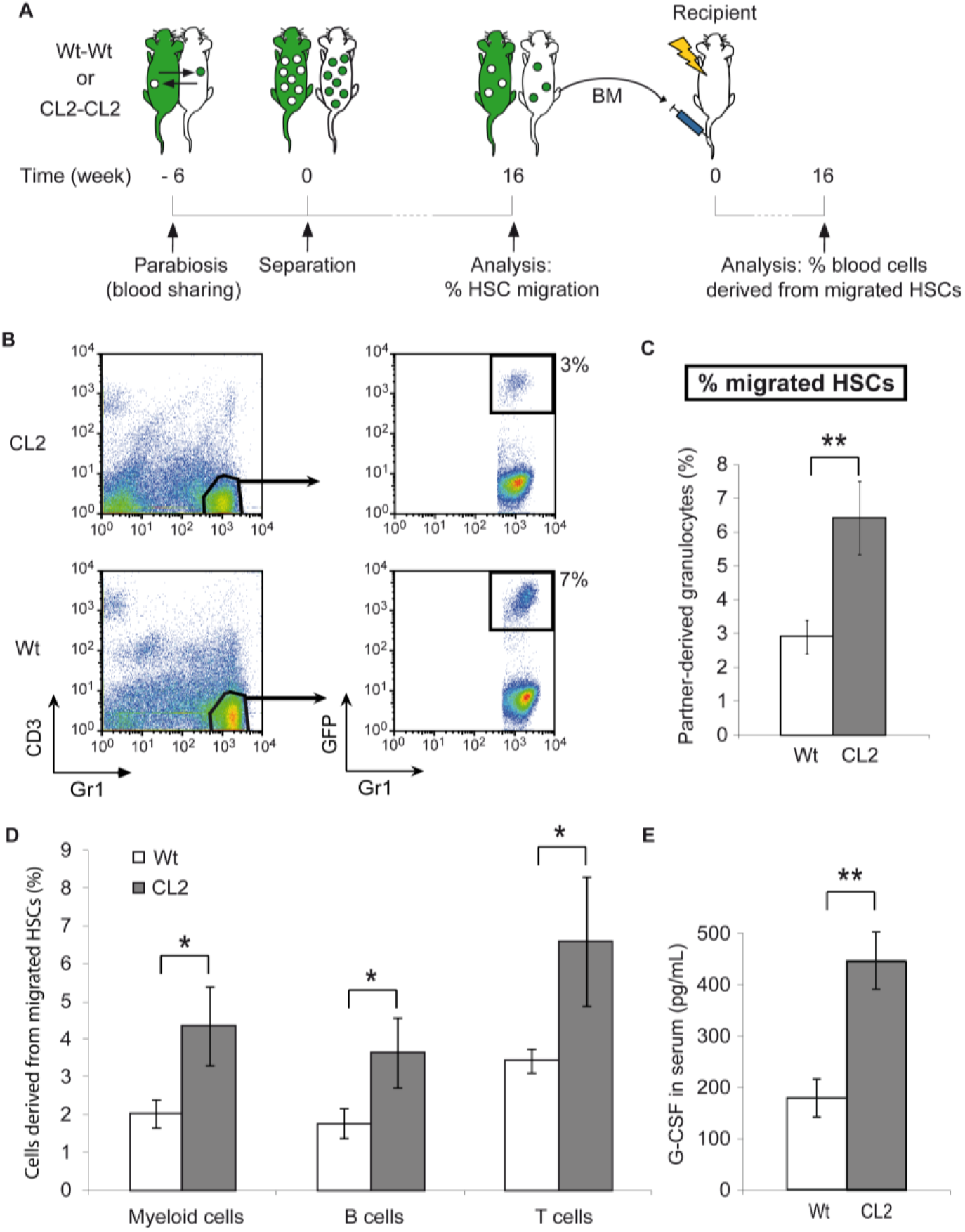
Higher HSC migration in CL2 mice. A. Schematic of the experimental design. B. Dot plots show a representative analysis of WT and CL2 BM. Cells were analyzed for Gr1, CD3 and GFP. BM of separated parabionts was harvested and the frequency of granulocytes derived from cross-engrafted HSCs was measured by flow cytometry. Gated regions designate granulocytes (left plots) and granulocytes issued from partner-derived HSCs (right plots). C. Graph shows the proportion of partner-derived granulocytes representative of the proportion of HSCs that have migrated and re-engrafted the partner’s BM, n=8. D. Whole BM cells from either WT or CL2 separated parabionts were transplanted into lethally irradiated primary recipients. The peripheral blood of recipient mice was harvested and among donor-derived cells (CD45.2^+^), the frequency of myeloid cells (Gr1^+^ and Mac1^+^), T cells (CD3^+^) and B cells (B220^+^) derived from cross-engrafted HSCs (GFP^+^) was measured by flow cytometry. Graph shows the frequency of differentiated cells issued from partner-derived HSCs at 4 months post-transplantation. E. Graph shows the G-CSF level in WT and CL2 serum. Each bar indicates the mean percent (± SEM) of at least 4 mice. *P<0.05 & **P<0.01.

### CL2 BM HSCs have impaired BM reconstitution capacity

We further wanted to know whether PTH-dependent activation of osteoblasts would influence the engraftment and BM repopulation properties of BM-HSCs. We therefore compared the ability of BM-HSCs from CL2 and WT mice to reconstitute WT-BM. To do so we performed a competitive long-term repopulation assay with total BM of CL2 and WT mice (Fig. 6A). As mentioned above, the mutant PPR is not expressed on HSCs due its collagenI-driven expression and therefore, it should not intrinsically influence HSC’s ability to reconstitute BM. Furthermore, since we found an equivalent proportion of HSCs in WT and CL2 mice, the long-term competitive repopulation assay should only reflect the ability of HSCs to home, engraft and repopulate BM. At 16 weeks post-transplantation, our data show that only 40% of CL2 HSCs participated in recipient’s BM repopulation indicating a CL2 BM significant competitive repopulation disadvantage compared to WT HSCs (Fig. 6B). These results suggest that PPR-dependent osteoblast activation negatively impacts on HSC properties to home, engraft and/or repopulate irradiated BM.

**Fig 6:**
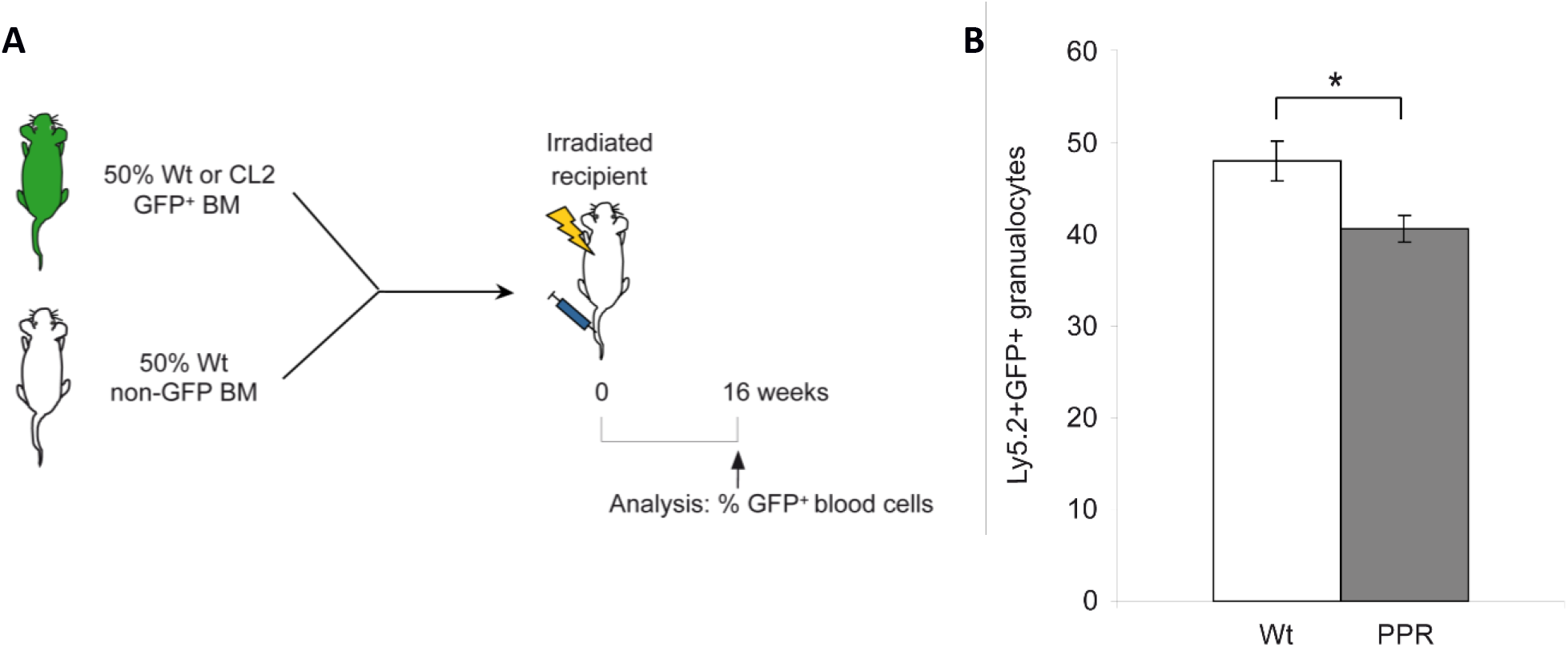
CL2 BM HSCs exhibit lower BM engraftment/repopulation ability in irradiated mice. A. Schematic of the experimental design. B. Graph representative of 4 independent competition assays. *P<0.05.

## Discussion

In this study we show that transgenic mice with PTH-dependent activated osteoblasts exhibit: 1) an increased hematopoietic progenitor cell rate without any change in HSC proportion in the BM, 2) low BM cellularity and HSC total numbers, 3) greater BM HSC migration into blood circulation and 4) a reduced ability of HSCs to repopulate irradiated BM.

The CL2 mice have previously been shown to exhibit PTH-dependent increased numbers of osteoblasts [19]. Calvi et al. have further demonstrated an increase of trabecular bone volume in CL2 mice due to higher osteoblast numbers and activity, resulting in remodeling of marrow cavity and a correlated increase in hematopoietic stem/progenitor cell numbers [2]. Our findings that the rate of hematopoietic progenitor cells is twice higher in CL2 BM than in the WT are in accordance with this study and with the idea that osteoblasts play a key role in hematopoiesis commitment [11]. However, our results indicate that the proportion of CL2 and WT BM HSCs are similar, which suggests that the PTH-dependent increase of osteoblasts only influences the rate of progenitors and has no effect on HSC frequency. These data are in agreement with several studies showing that osteoblastic cell depletion or strontium increased osteoblasts has no significant impact on HSC rate [16–17, 27]. Furthermore, recent works based on deep confocal imaging and specifically GFP-labeled HSCs demonstrate that HSCs are mainly located in perisinusoidal niches [8]. Our results are in accordance with the notion that osteoblasts might be further involved in more committed progenitors. Indeed, several studies have demonstrated that osteoblasts are further involved in hematopoiesis and support B lymphocyte maintenance and differentiation [16, 28–30]. Therefore, our data support the idea that the osteoblastic niche would further be a progenitor niche than an HSC niche [11].

The activation of osteoblasts in CL2 BM is the consequence of a constitutively active form of the PTH receptor, mimicking PTH anabolic effect specifically on osteoblasts. In our study, we found increased HSC migration into blood circulation of CL2 mice. While the main mobilizing molecule used in clinics to induce hematopoietic stem/progenitor cell mobilization is G-CSF, PTH was also shown to display the same properties of mobilization [20, 31–33]. More recent works demonstrated that PTH treatment could increase G-CSF-induced stem/progenitor cell migration into blood circulation [21]. The same study also shows that secondary recipients transplanted with BM from primary recipients treated with PTH get higher BM repopulation. In our study we further demonstrate that without G-CSF treatment or BM irradiation, CL2 mice exhibit higher LT-HSC migration into blood circulation. While mechanisms influencing HSC migration are poorly understood, it is well known that HSCs enter and exit the BM via endothelial cell made blood vessels. Furthermore, studies based on living imaging methods reveal that the BM of CL2 mice is more vascularized than WT BM, which could partly explain the increased HSC migration into CL2 blood circulation [34]. Furthermore, our results showing higher rates of G-CSF in CL2 mice suggest that HSCs may also be mobilized by this factor. These results are in accordance with some works that also measured higher G-CSF levels in peripheral blood after PTH treatment [20]. PTH has been proposed as new mobilizating agent for treatment of hematological disorders. Our data demonstrating that PTH induces LT-HSC mobilization into blood circulation without the need for myeloablation conditions or G-CSF treatment reinforce previous ideas that PTH could be used as an alternative HSC mobilizating agent in clinics. However it is important to investigate whether HSCs keep their self-renewal and differentiation properties after a PTH treatment.

The property of a mobilizating agent is to mobilize BM stem/progenitor cells into blood circulation. However, in order to cure hematological disorder patients, mobilized cells have to keep their BM repopulation properties. In our study, the BM competitive repopulation experiments reveal that BM HSCs coming from a CL2 donor mouse fail to efficiently reconstitute the BM of irradiated WT recipients. Our data suggest that PTH-dependent activation of osteoblasts negatively influences HSC capacities to reconstitute irradiated BM. The reconstitution of BM is a multistep process including homing, engraftment and HSC proliferation and differentiation into all blood cell lineages [11, 35–37]. It is well known that adhesion molecules play a key role in HSC homing and BM engraftment. Indeed, several studies have demonstrated the role of α4 integrins in HSC homing and BM repopulation [36, 38]. Similar to our results HSCs were found to display a repopulation defect when lacking α4 integrins [39]. The chemokine receptor CXCR4 expressed on HSCs is also involved in HSC homing into irradiated BM [40]. Therefore, these studies altogether with our findings that CL2 HSCs have impaired BM reconstitution suggest that PTH-dependent activation of osteoblasts may influence the expression of such adhesion molecules and therefore that CL2-HSCs may lack on their surface the adhesion molecules required for homing/engraftment. Furthermore, several mobilization factor treatments such as with G-CSF led to reduced HSC homing due to a cleavage of the CXCR4 N-terminus [41]. In line with these findings, we found increased G-CSF levels in CL2 mice, which could explain the HSC defect in BM reconstitution. Another explanation for the poor BM reconstitution outcome when osteoblasts are exposed to PTH could be the heterogeneity among HSCs. Indeed, Oguro et al. by using the SLAM family markers have identified different HSC subpopulations [7]. Furthermore, recent studies based on single cell phenotyping have highlighted HSC heterogeneity in the BM [42]. Therefore, since CL2 mice display higher HSC migration into blood circulation, it could be possible that HSCs remaining in the BM constitute one HSC subtype with lower BM repopulation properties. Furthermore, it was proposed that activated and quiescent HSCs would reside in different niches [11]. The CL2 mice display an increase of trabercular bone [19]. Therefore, this bone structure remodeling could have modified the different niche distribution and impacted on HSC properties. Another possibility could be that the defect of CL2-HSCs in BM competition repopulating assay would reflect a loss in their proliferation or differentiation properties. Human HSCs located in the long bone area have been shown to display lower regenerative and self-renewal capacities than HSCs located next to trabecular bones [43]. Therefore, since CL2 mice exhibit higher trabecular bones, CL2 HSCs should be more efficient in regenerating BM, which is in contradiction with our findings. An explanation could be that CL2 HSCs might not locate in the trabecular bone area. Further analyzes on HSC localization in CL2 BM will allow to better understand the role of HSC microenvironment on their trafficking, engraftment and BM repopulation abilities. Moreover, it would be interesting to sequentially examine each step leading to irradiated BM repopulation to determine the role of PTH in homing, engraftment and HSC proliferation and differentiation. Another important point would be to study the PTH influence on the expression of HSC surface proteins involved in the BM repopulation process.

### Conclusion

In summary, our study shows that the PTH dependent activation of osteoblasts influences BM HSC properties by increasing their mobilization into blood circulation and by reducing their capacity to efficiently reconstitute ablated BM. These data shed some light on the role of PTH and osteoblasts on HSCs, which would be useful for future therapies in clinics to improve hematological disorders.

## Acknowledgements

This work was supported by a CIHR grant MOP 160678 to F.M.R. and Y.E. was supported by a postdoctoral fellowship from la Fondation pour la Recherche Medicale (FRM).

We thank Dr D. Scadden and Dr E. Schipani for the kind gift of CL2 mice. We thank A. Johnson for help with flow cytometry, M. Williams for antibody production, J.H. Kang and B. Paylor, for technical help and S. Sekulovic for CRU statistical analysis assistance. This work was supported by a CIHR grant MOP 160678 to F.M.R. F.M.R. holds a Canada Research Chair in Regenerative Medicine. Y.E. was supported by a postdoctoral fellowship from la Fondation pour la Recherche Medicale (FRM).

